# Nanoscale molecular architecture controls calcium diffusion and ER replenishment in dendritic spines

**DOI:** 10.1101/2021.06.08.447402

**Authors:** Kanishka Basnayake, David Mazaud, Lilia Kushnireva, Alexis Bemelmans, Nathalie Rouach, Eduard Korkotian, David Holcman

**Affiliations:** Computational Biology and Applied Mathematics, Institut de Biologie de l’É cole Normale Supérieure-PSL, Paris, France; Neuroglial Interactions in Cerebral Physiology, Center for Interdisciplinary Research in Biology, Collège de France. CNRS UMR 7241, INSERM U1050, Labex Memolife, PSL Research University, Paris, France; Faculty of Biology, Perm State University, Perm, Russia; Commissariat à l’Energie Atomique et aux Energies Alternatives, Département de la Recherche Fondamentale, Institut de biologie François Jacob, Molecular Imaging Research Center and Centre National de la Recherche Scientifique UMR9199, Université Paris-Sud, Neurodegenerative Diseases Laboratory, Fontenay-aux-Roses, France; Department of Neurobiology, Weizmann Institute of Science, Rehovot, Israel; Churchill College and the Department of Applied Mathematics and Theoretical Physics, University of Cambridge, Cambridge, United Kingdom

## Abstract

Dendritic spines are critical components of the neuronal synapse as they receive and transform the synaptic input into a succession of biochemical events regulated by calcium signaling. The spine apparatus (SA), an extension of smooth endoplasmic reticulum (ER), regulates slow and fast calcium dynamics in spines. Calcium release events from SA result in a rapid depletion of calcium ion reservoir, yet the next cycle of signaling requires replenishment of SA calcium stores. How dendritic spines achieve this without triggering calcium release remains unclear. Using computational modeling, calcium and STED super-resolution imaging, we showed that the refilling of calcium-deprived SA involves store-operated calcium entry during spontaneous calcium transients in spine heads. We identified two main conditions that guarantee SA replenishment without depletion: (1) a small amplitude and slow timescale for calcium influx, and (2) a close proximity between SA and plasma membranes. Thereby, molecular nano-organization creates the conditions for a clear separation between SA replenishment and depletion. We further conclude that the nanoscale organization of SA receptors underlies the specificity of calcium dynamics patterns during the induction of long-term synaptic changes.

## Introduction

Dendritic spines are cellular protrusions found on the surface of dendrites that function as a postsynaptic component of the neuronal synapse. Morphologically, spines commonly include two distinct features, a spine head and a spine neck, with the head engaging the presynaptic axon and the neck connecting the head with the postsynaptic dendrite. The size and shape of a spine (as well as the number of spines on a dendrite) are dynamic, plastic, and change in response to repeated synaptic activity, which directly impacts synaptic plasticity and is therefore critical for processes such as learning and memory (*1–4*). Moreover, changes in shape, distribution, loss as well as gain of dendritic spines has been associated with a range of human diseases, although the mechanisms remain incompletely understood (*5, 6*). Functionally, dendritic spines are sites of intense biochemical activity, whereby signals received from the synapse via glutamate receptors (such as NMDARs and AMPARs) lead to influx of the main second messenger, calcium ions (Ca^2+^). Although a single receptor activation event produces transient calcium influx, repeated activation of the receptor leads to rapid build-up of calcium concentration, resulting in calcium binding to buffers such as calmodulin (CaM), a calcium sensor protein, and subsequent biochemical activation of calmodulin-dependent protein kinase II (CaMKII) and subsequent downstream signaling (*7*). In general, high calcium concentrations in spines are associated with long term-potentiation (LTP), while low calcium is associated with long term depression (LTD) (*8*). Thus, calcium dynamics in dendritic spines emerged as a key mechanism for the induction of synaptic plasticity in neurons. Calcium concentration in dendritic spines is tightly regulated (*9–11*), by the amplitude and timing of the influx through the receptors, binding and unbinding with proteins, buffers and pumps, and also by organelles that could sequester or release calcium, such as endoplasmic reticulum (ER) and an extension of smooth ER called spine apparatus (SA) that spans spine head and neck (*3, 12–14*).

The SA is involved in multiple signaling functions such as calcium regulation, protein synthesis and cell apoptosis, and the absence of the SA results in a reduction of hippocampal long-term potentiation in the CA1 region and an impairment of spatial learning (*15*). Additionally, SA facilitates calcium-induced calcium release (CICR) process through which calcium ions that entered dendritic spine due to synaptic activity cause release of additional calcium from intracellular calcium stores. The timing of CICR activation, on the order of a few tens of milliseconds, is governed by the fastest calcium ions arriving by diffusion at a ryanodine receptor (RyR) present mostly as clusters at the base of spines (*16*), and also involves sarco/endoplasmic reticulum Ca^2+^-ATPase (SERCA) (*17*). Therefore, calcium concentration increase in a spine head due to a synaptic stimulation is followed by a pronounced depletion of SA calcium reservoir within less than tens of milliseconds due to this avalanche phenomenon at the base of the spine. Whereas CICR calcium dynamics and flux in spines have been extensively studied, mechanisms that govern calcium store replenishment in SA is still imperfectly understood.

Recently, store-operated calcium entry (SOCE), mediated by STIM1-ORAI1 channel complex, has been implicated as important for the replenishment process in SA (*18, 19*). Calcium concentration inside the SA is sensed by ER-membrane anchored STIM1 regulatory protein that interacts with ORAI1 channel present at the plasma membrane (*20*). As the first step, the ORAI1 channels pump calcium into cytoplasm, followed by calcium entrance into the SA through SERCA pumps mostly located in the SA head (*16*). However, it remains unclear how SOCE could function to replenish calcium stores without triggering CICR activation. Here we investigate the release and replenishment pathways that are both activated by calcium but could operate without interfering with each other. Resolving this enigma is crucial, not only to determine the computational power of dendritic spines based on calcium signaling, but also for characterizing calcium dynamics underlying the induction of LTP and LTD. To study these processes under different calcium influx conditions (large and small, fast and slow), we employed a combination of calcium imaging, computational modeling and simulations, and STED microscopy. Our stochastic model predicted and our measurements confirmed that SA-amplified CICR occurs only under strong and fast calcium influx conditions, where as slow and small amplitude calcium influx triggers SA replenishment via STIM1-ORAI1 pathway and SERCA pumps located proximal to the plasma membrane. Furthermore, we also examined using numerical simulations calcium dynamics at the base of spines, during LTP and LTD and observed that the SA depletion timescale varies between LTP and LTD, resulting in a strong difference in the calcium levels at the base, which may determine the spine’s fate towards the direction of either enhancement or depression of synaptic efficacy. Taken together, our study reveals that nanoscale molecular architecture plays a critical role in controlling calcium diffusion and ER replenishment. Moreover, contrary to the view that calcium concentration inside the whole spine dictates potentiation or depression, the work presented here suggests that calcium concentration at the spine base is the main determinant of LTP versus LTD induction.

## Results

### SOCE is associated with SA replenishment but not depletion

To investigate the mechanism of SA replenishment, following the methods developed in (*21*) and (*22*), we blocked synaptic activity (calcium voltage channels (CaV) and synaptic inputs using APV/DNQX/TTX) in cultured hippocampal neurons co-transferred with blue fluorescent protein and synaptopodin (SP), an actin-associated protein found in SA (*12*). The only known source of calcium in these conditions remains the SOCE mechanism associated with the STIM1-ORAI1 pathway (*20*). In order to monitor calcium fluctuations in spines containing SA, we used Fluo-4, a high-affinity calcium sensor (See also Materials and Methods). With this setup, we observed fluctuations that were restricted to spines and were not present in dendrites (Fig. 1A-B top panels). Additionally, these calcium activity patterns associated to the STIM1-ORAI1 complex were much slower (on the order of seconds) and exhibited smaller amplitudes compared to the ones triggered by synaptic inputs (on the order of a few hundred milliseconds: Fig. S1 and S2).

**Figure 1:**
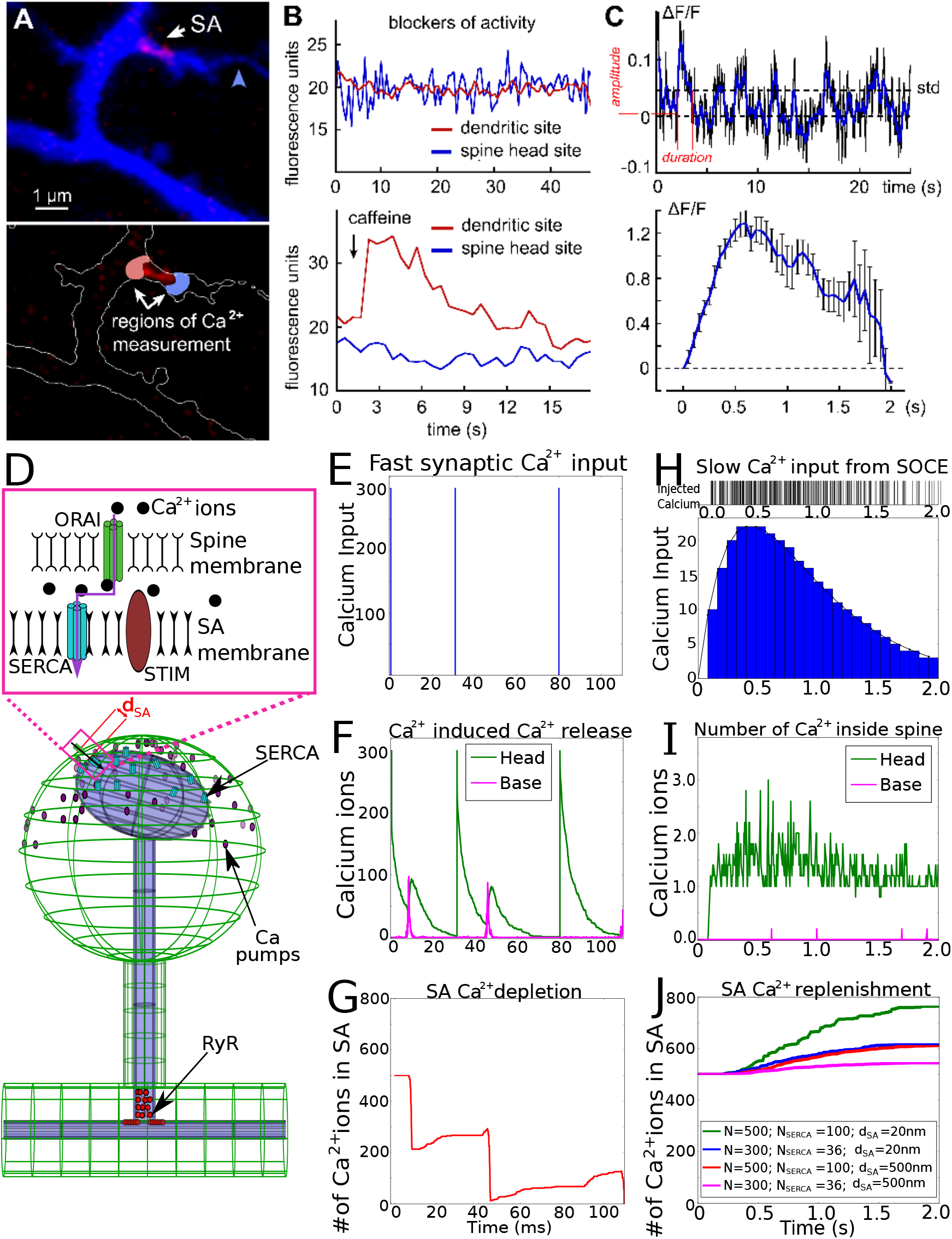
Refilling or depleting the SA in dendritic spines with slow versus fast calcium transients. **(A).** Top: Blue fluorescent protein (blue) and SP (red) co-transfected in hippocampal neurons, cultured 3 weeks and loaded with the Fluo2 high affinity calcium sensor. A large SP-positive spine attached to an axon (blue arrowhead) and several SP-negative spines can be seen. Bottom: Same region, with white contour and two regions of interest: in front of the SP-labelled SA (blue) and behind SP (red) are shown. Synaptic activity was blocked using tetrodotoxin (1*μ*M), APV (50*μ*M) and DNQX (20*μ*M). Following a caffeine addition of 5 mM, release of calcium is observed towards the base of the spine, but not in the spine head. **(B).** Top: spontaneous calcium activity due to SOCE, only located in the spine head. Bottom: time course of calcium activity in the head vs base following caffeine addition. Both are typical, single realisations of the experiments. **(C).** Top: segmented recording of spontaneous calcium activity in the spine head. Bottom: overlapped average of the fluctuation segments larger than a pre-defined threshold of one standard deviation (std=0.0457) in the example trial shown in the top panel. Bottom panel shows the average of 18 such sequences. **(D).** Schematic representation of the SOCE regulators in the spine. In the computational simulations, calcium ion input is generated at the top of the spine head. Ions diffuse inside the domain (green) and eventually reach the bottom of the spine where they are removed from the simulations. Calcium pumps on the spine head, SERCA pumps on the SA head and RyRs at the base could capture the diffusing ions (See Materials and Methods for the modeling of each channel type). **(E).** Simulated synaptic input with *N*=300 calcium ions instantaneously placed in the spine head. The first input is at t=0, and we repeat each time when the calcium in the spine decays to zero (green curve: panel F). Thus the second and the third inputs were at t=30.9s and t=80.1s. **(F).** Typical simulation result shows a sharp increase of calcium due to CICR occurring at the base, a few milliseconds after each calcium injection into the head. Calcium decay in the head is due to the uptake through SERCA pumps, the diffusion towards the base and the absorption by pumps. Smaller second peaks (about 10ms following each input spike) manifest due to calcium ions “generated” by CICR at the base and diffuse into spine head. **(G).** The number of calcium ions in the SA reservoir decreases systematically after each stimulation due to the CICR at the base. After the third release, all 500 ions initially in the SA are depleted. **(H).** We approximate slow calcium activity in spines as a difference of exponentials *I*(*t*) = *Q.*[e^*−*2.3*.t*^ e^*−*2.31*.t*^] (black curve). We consider a total duration of 2s (which accounts for 94.5% of the distribution) and discretize this by dividing into 25 equal bins with integer values (histogram), and normalized such that the total number of calcium in 2s is *N* (we choose either *N*=300 or 500). This approximation has *R*^2^ = 0.9986. Single calcium ion injections times then follow a uniform random distribution according to the number of ions corresponding to each time bin. (Barcode representation: top panel). **(I).** Number of calcium ions in the spine head and the arrivals at the base of the spine (average over 5 trials with *N*=300). Maximum calcium arrival at the base was about 1 ion, and no calcium release events occur from RyRs. **(J).** Total number of calcium ions in the SA for different distances (*d*_SA_) between spine head and SA, and different number of SERCA pumps *N*_SERCA_. In all cases, the SA calcium level is increasing gradually via SOCE refilling throughout the 2s duration of the calcium input because SA is not depleted by CICR events.

To confirm that the SOCE pathway leads to calcium accumulation in the SA, we depleted the SA calcium stores with caffeine (*12*) (Fig. 1B bottom panel) and found an asymmetric calcium release, mostly toward the base of the spine. This result is in agreement with the timescale of CICR activation and the underlying heterogeneous distribution of RyRs that are mostly located at the base of the spine (*16*). To further study calcium transients and develop numerical simulations, we needed to obtain a stereotype response. For that goal, we segmented the calcium time series, which was recorded in the spine head over a timescale of a few minutes. We defined a threshold which is one standard deviation (std) to differentiate between calcium transient and background fluctuations (std ≈ 0.0457 in Fig. 1C top panel). We collected events and averaged them (Fig. 1C bottom panel) resulting in a stereotype response that we fitted with a difference of two exponentials (Fig. S3). The calcium concentration in the spine head has a correlation time of approximately 1s, as revealed by the autocorrelation function (Fig. S4), compatible with previous analysis on calcium transients in (*23*). We then used this fit to determine the conditions that favor calcium accumulation in the SA by developing stochastic computational simulations for calcium diffusion inside a dendritic spine (Fig. 1D). For calcium inputs to spine head, we used two distinct conditions: (a) a fast entry from synaptic inputs through NMDA receptors, which we modeled by injecting calcium into the spine head instantaneously; and (b) a slow calcium entry by slow injection, which was modeled to mimic what happens during SOCE activity (as observed in Fig. 1C-lower panel). Our results revealed that:

A. fast synaptic input is associated with store depletion events through RyR-induced CICR. In the numerical simulations (Fig. 1E), we verified this prediction by injecting 300 ions into the spine head to mimic fast synaptic inputs, then waiting until the spine head was completely depleted of calcium, before injecting another batch of 300 ions. In all, we thus simulated three injections, at t=0s, 30.9s and 80.1s and observed a sharp spike in calcium concentration at the spine base within few milliseconds of the injection, due to CICR (Fig. 1E-G). This was accompanied by the loss of calcium in the head through calcium pumps or diffusion to the dendrite (Fig. 1E-G). After each injection, we also observed a much smaller second calcium peak after a few milliseconds in the head, due to diffusion of CICR-released calcium from the base to the head region. When we started with 500 ions in the SA, after three stimulations, calcium ions in the SA became depleted due to CICR as predicted under these conditions (*24*). Note that the small increases in between the depletions (Fig. 1G) are due to uptake of ions into SA through SERCA pumps.
B. In contrast, when we injected ions at a slow rate into the spine head compared to synaptic inputs (Fig. 1H) by accounting for the response above threshold std (as observed in Fig. 1C bottom panel), we found that although the number of calcium ions in the head was much reduced compared to levels under synaptic input conditions (Fig. 1I), the SA was refilling by calcium without triggering CICR (Fig. 1J). For this mechanism to occur, we positioned SERCA and ORAI1 in close proximity in the spine head and we hypothesized that the distance *d*_SA_ between the SA and the plasma membrane is in the order of a few tens of nanometers.

We explored the impact of *d*_SA_ on SA refilling by changing the distance from 20nm to 500nm (Fig. 1J). We found that the reduction of flux due to the increased distance can be compensated by increasing the number of SERCA pumps from 36 to 100 or by increasing the calcium input from *N*=300 to 500 ions. Therefore, these results suggest that the distance *d*_SA_, the number of SERCA pumps as well as the slow influx of calcium are key parameters favoring SA calcium refilling inside the spine.

To further characterize the distribution of calcium fluxes in the SA, (extrusion from the head or the arrival at the base of the spine), we followed the fate of each ion during numerical simulations. We simulated 300 injected ions each time over a 2s duration, keeping the distance between the membrane constant at 20nm (similar to Fig. 1H). In an ensemble of 210 trials we simulated, we analysed the 161 in which a CICR did not occur. We observed that most ions (236/300) were bound to calcium extrusion pumps, while the number of ions that refilled the SA through SERCA pumps was 46/300. Around 14/300 remained in the spine domain or stayed as single-occupants in SERCA or RyRs without triggering an opening within the simulated 2s; only the remaining 4/300 ions reached the base and disappeared from the absorbing boundary. Therefore, under these conditions, calcium concentration at the base of the spine remained too low to trigger a CICR, safely eliminating the possibility of triggering a depletion.

To characterize the spatio-temporal conditions of calcium dynamics leading to SA replenishment, we also developed a mean-field computational model (Section S 4), accounting for the mean number of calcium ions *m_ca_* in the spine, the fraction *n*_1_ of RyRs bound by one calcium ion and the probability *p*_2_ to trigger CICR by activating at least one RyR when two calcium ions are bound. The complete system of equations are:

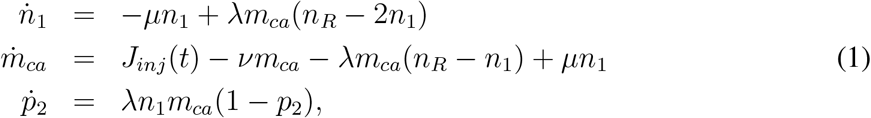

where *μ* is the unbinding rate of calcium to RyR, *ν* is the calcium extrusion rate from a spine, and *λ* is the forward rate of calcium to RyR. To study the fast synaptic inputs and SOCE, we used the two different initial conditions: (1) ions were injected instantaneously, and modeled with a Dirac delta function at *t* = 0 (or equivalently with the initial condition *m_ca_*(0) = *N*_0_*, n*_1_(0) = 0*, p*_2_(0) = 0); (2) a slow injection rate, modeled by 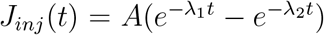 (Fitted in Fig. S3), with the initial conditions of *m_ca_*(0) = 0*, n*_1_(0) = 0 and *p*_2_(0) = 0. The probability to activate RyR by two ions is

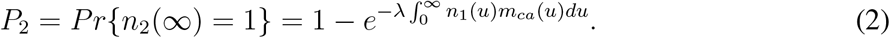

We evaluated numerically the activation probability *P*_2_ (Fig. S8), and confirmed its agreement with the results of the stochastic simulations Fig. 1: even with a small (<500) number of ions, an instantaneous calcium influx *J_inj_* (*t*) leads to a higher probability of SA calcium release, whereas a slower influx with such a number leads to CICR events at a much low probability. The solution of equations 1 (Fig. S5 and the Sections S4.2-S4.3) shows how the probability *P*_2_ depends on the number *N* in fast synaptic versus the amplitude *A* in the slow SOCE inputs. Therefore, both our mean-field model and the stochastic simulations strongly indicate that a slow calcium transient from the STIM1-ORAI1 pathway does not induce calcium release at the base of a dendritic spine, but leads to SA refilling. Thus we conclude that the biophysical conditions for SA replenishment and depletion are well separated (Fig. S6). Collectively, our calcium imaging-based observations and our computational modeling suggest that SOCE plays role in replenishment exclusively, and that spatial co-localization (proximity) of key molecular regulators in this process (SERCA and ORAI1) is critical for ensuring fidelity of replenishment while not triggering CICR.

### Calcium redistribution in dendritic spines: replenishment without release

To test the predictions generated using our computational modeling of SA replenishment, we explored the conditions for calcium replenishment in dendritic spines containing an SA using calcium imaging of hippocampal cultured neurons. To identify spines containing SA, we used SP as a marker for SA (red puncta in Fig. 2A&B). To demonstrate that the SA is refilled in the absence of any activity, we blocked both voltage-gated channels and glutamatergic receptors by adding tetrodotoxin (TTX, 1*μ*M), APV (50*μ*M) and DNQX (20*μ*M). Indeed, in the absence of extracellular calcium, spines do not show any calcium transients regardless of whether they contain SA or not (Fig. 2C1-C2). By blocking the SOCE pathway (Fig. S1), we confirmed that STIM1-ORAI1 is responsible for SA calcium replenishment. Moreover, we observed that the replenishment of calcium in SA through SOCE increases with duration of refilling (Fig. S2). In contrast, when presented with extracellular calcium, significant transients occur only in the heads of SP+ spines (Fig. 2D1-D2-D3). Both the amplitudes (2.2 vs 1.54 a.u.) and the durations (1.4s vs 0.8s) are significantly larger in SP+ spines compared to SP-ones (Fig. 2 D4). In addition, the frequencies of activation (Fig. 2D5) were clearly higher in the spine head of SP+ (grey) compared to SP-spines (purple) or dendrites (black). Therefore we conclude that SOCE occurs preferentially in the heads of SP+ spines and requires the presence of SA.

**Figure 2:**
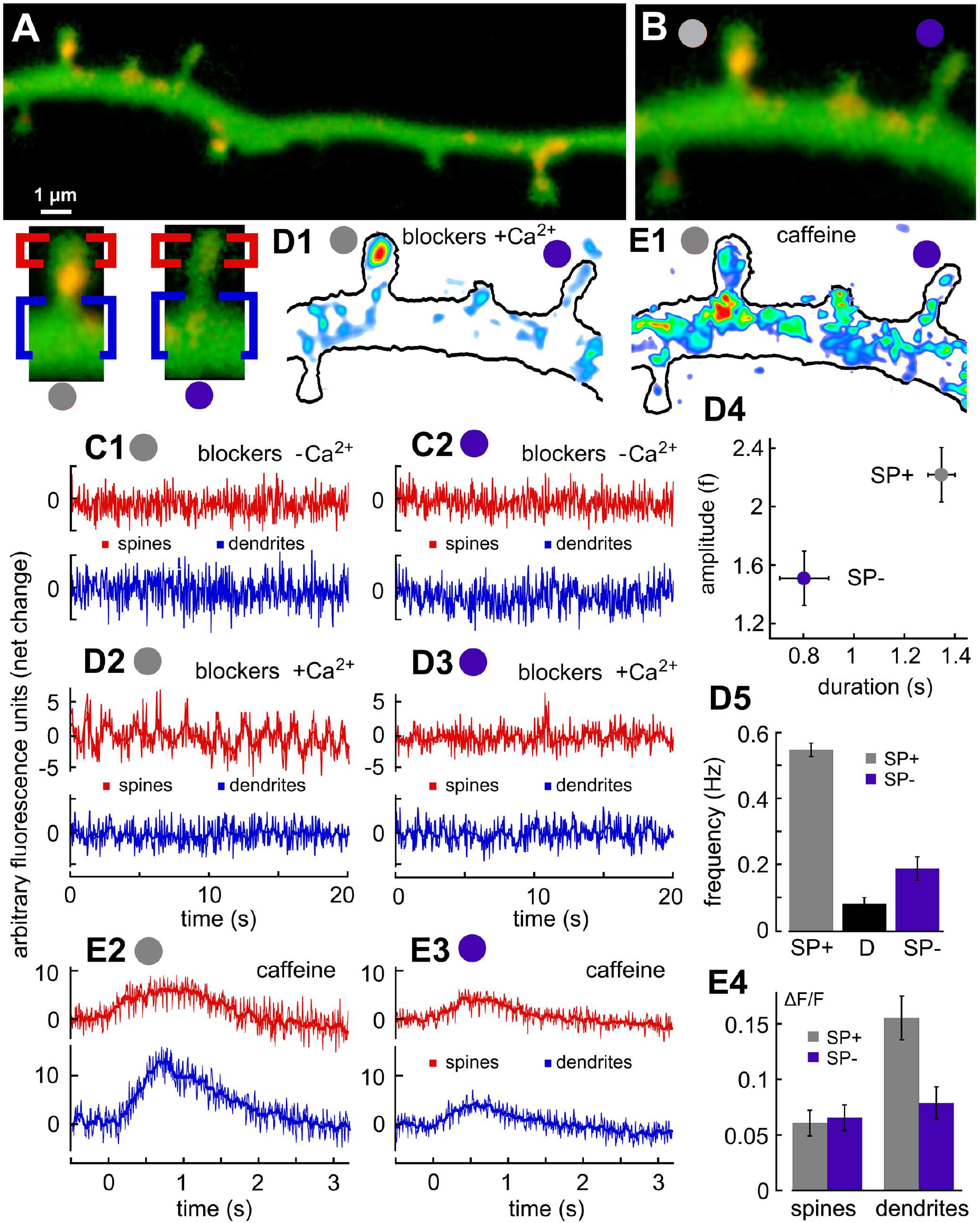
Store-operated calcium loading into synaptopodin-associated local calcium storage. **(A)** Dendrite of rat hippocampal cultured neuron, transfected with synaptopodin (SP) (red puncta) and loaded with high affinity calcium sensor Fluo-2. **(B)** Magnification of the dendritic fragment shown in (A) with two dendritic spines of approximately same length of 1.2 *μm*). SP+ spine (we use the gray circles to show SP+) is on the left and SP-(violet circle) is on the right. Same spines are shown below panel A, on the left. Here, we label the region of head edge with red, while the basal edge close to dendritic shafts are shown in blue. **(C1)** (SP+) and **(C2)** (SP-) calcium recordings from these two regions of both spines in calcium-free medium in the presence of following activity blockers: APV (50 *μ*M), DNQX (20 *μ*M) and TTX (1 *μ*M). No activity is recorded in any compartment of both spine types. After applying these conditions for 15 mins, calcium stores became partly depleted and store-operated calcium entry mechanism is initiated (not shown). **(D2)** and **(D3)** Same regions are recorded with activity blockers but under the presence of extracellular calcium (2 mM). Low amplitude calcium fluctuations are regularly seen in the head of SP+ spine but rarely seen in SP- spine or in the adjacent dendrites. Example of SP-head-located transient is shown on **(D1)** (background subtracted, spectral color code is used to show the calcium level: low calcium level is represented by blue/cyan while higher calcium levels are from red>yellow>green). **(D4)** Overall averaged amplitudes and durations of calcium fluctuation were significantly larger in SP+ (gray dot) than in SP- spines (violet dot) (n =16 for both groups, p<0.0001, t-test). **(D5)** Frequency of calcium fluctuations was also significantly higher in SP+ group (0.5 Hz, gray column) than in adjacent dendrites (0.1 Hz, black column) or SP-group (0.2 Hz, violet column) (n=16, p<0.0001, ANOVA). **(E2) and (E3)** Same regions following caffeine bath application (5 mM) which releases calcium from internal stores. Calcium release occurs towards the shaft and only in SP+ spines. Examples of caffeine-induced calcium transient is shown on **(E1)** (background level is subtracted, same color code as D1). **(E4)** Caffeine responses in SP+ heads, SP- dendrites as well as SP-heads were approximately the same, but significantly lower than the stronger SP+ dendritic site responses (n= 16 for all groups, *p <* 0.001, ANOVA).

Interestingly, calcium entry did not lead to any calcium transient at the bases of both SP+ and SP-spines, confirming our predictions (showed in Fig. 1) that a slow calcium entry leading to SOCE does not activate RyRs. However in order to confirm that RyRs were functional, we activated them using caffeine. This activation led to significant transient increases of calcium at the base of SP+ spines only, where RyRs are mostly located (Fig. 2 panels E1-E3). Indeed in these experiments, caffeine is present in the extracellular medium and trigger CICR with a slower time scale compared our simulations where calcium ions already present in the spine head. In summary, these results confirm the initial theoretical predictions that in the absence of synaptic activity, calcium ions enter through SOCE and are stored inside the SA.

### STED microscopy reveals the colocalization of ORAI1 in plasma membrane and SERCA pumps in spines with a SA

Our stochastic simulations predicted that SOCE is the main source of calcium ions for the SA calcium replenishment process. We thus hypothesized that ORAI1, which allow slow calcium entry from the plasma membrane should be closely colocalized with SERCA pumps that refill the SA. In addition, we predicted that this colocalization should predominately occur in the heads of spines containing SA. To determine the association between SERCA and ORAI1 localization, we used super-resolution STED imaging of immunostained hippocampal tissues (see methods).

We confirmed that the colocalization frequency of ORAI1 and SERCA3 is significantly higher when SP is present (ORAI1+SERCA3+SP: 88.9±0.73%, n=19, p<0.001) than when SP is absent (ORAI1+SERCA3: 0.99±0.24%, n=19, p<0.001). The colocalization of SP with ORAI1 or SERCA3 only is weak (ORAI1+SP: 7.75±0.61%, n=19, p<0.001; SERCA3+SP: 0.45±0.16%, n=16, p<0.001) similar to spines with SP alone (1.88±0.33%, n=19, p<0.001). Moreover, the average co-localization distance in SP+ heads is around 100nm, with a significant population of SERCA and ORAI1 localized as closely as 30nm, while their average distance is more than doubled to around 200nm in the neck or at the base of a spine (Fig. 3C-D). To conclude, these results reveal the short distance between SERCA and ORAI1 in spine heads containing spine apparatus, confirming the predictions of the stochastic simulations about SOCE-associated SA refilling.

**Figure 3:**
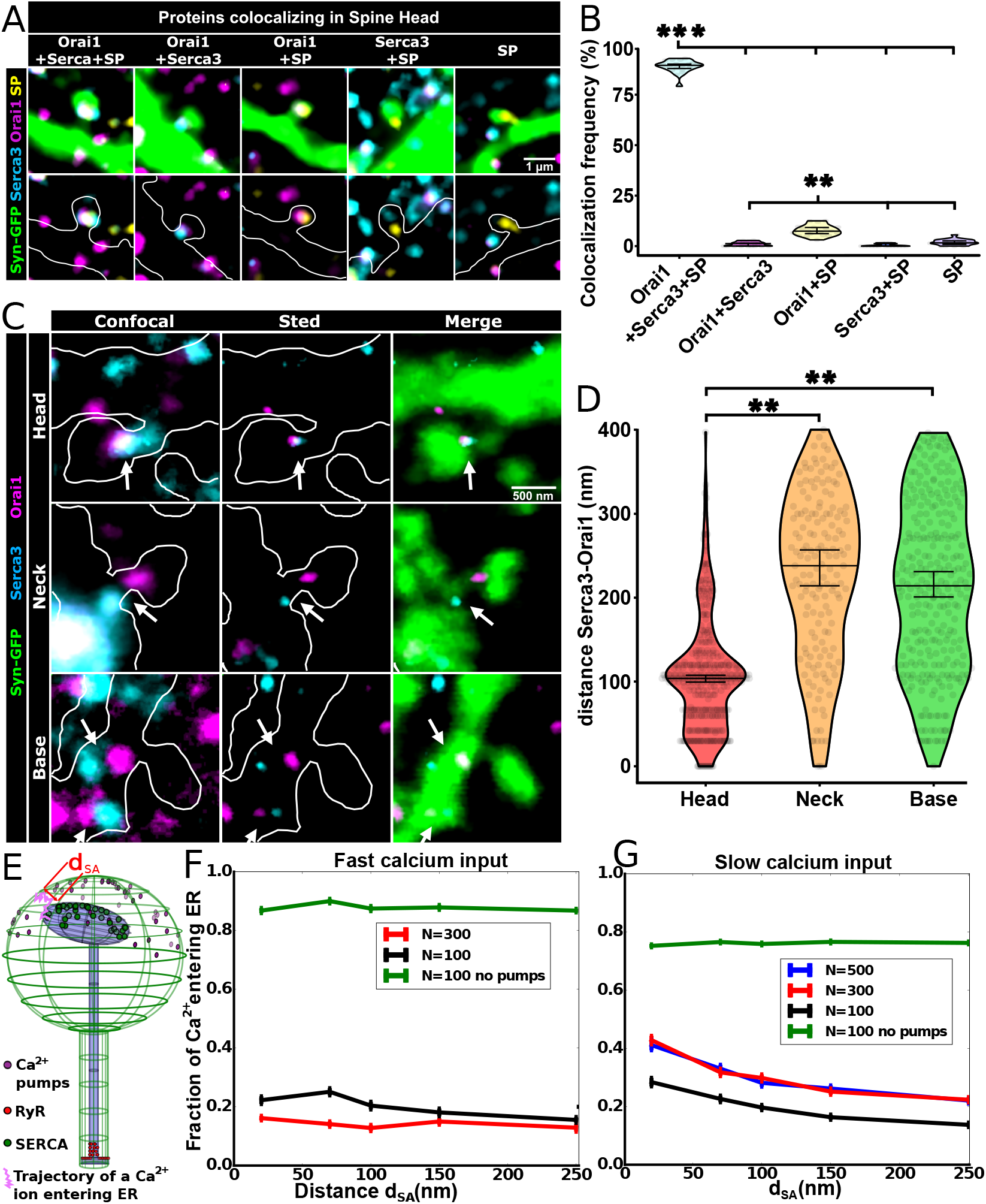
Statistics of SERCA-ORAI1 distributions and consequences. (A–D) Distance between SERCA3 and ORAI1 puncta in dendritic spines of hippocampal neurons from adult mouse brain slices depends on their localization within the spine and the presence of SP puncta. **(A)** GFP-labelled mouse hippocampal neurons (green on the left panels and white contours below) and immunostained for ORAI1 (magenta), SERCA3 (cyan) and SP (yellow). Representative images show the five conditions analyzed in panel B. Scale bar: 1 *μ*m **(B)** Quantification of the colocalization frequency between ORAI1, SERCA3 and SP proteins. The colocalization frequency of ORAI1 and SERCA3 in spine heads is significantly higher when SP i s present (88.9±0.73%, n=19, p<0.001) than when SP is absent (0.99±0.24%, n=19). Between 279 and 364 spine heads were analyzed per slice with four slices analyzed. **(C)** GFP-labelled mouse hippocampal neurons (shown in green on the right panels and delimited with white contours on the left panels) and immunostained for ORAI1 (magenta in STED and red in confocal panels) and SERCA3 (cyan in STED and blue in confocal panels). Representative images show the three situations analyzed in panel D: dots localized in the head, neck or base of the dendritic spine. Scale bar: 500 nm. **(D)** Quantification of the distances between SERCA3 and ORAI1 in dendritic spine head, neck or base. SERCA3 and ORAI1 are significantly closer to each other in the head (106±4 nm, n=378) than in the other parts of the spine (neck: 221±14 nm, n=159, p=0.0067; base: 210±8 nm, n=290, p=0.0034). Statistical analysis was performed with a oneway ANOVA test followed by Tukey’s multiple comparisons test. **(E)** Schematic of the calcium regulators in the simulated spine model showing the distance *d*_SA_ between the plasma and SA membranes. **(F)** Normalized fractions of calcium entering SA instantaneously to the head for various *N* with (black and red) and without (green) 50 calcium pumps in the spine head. Out of 100 trials, we selected the ones that did not activate a RyR until there were no more ions to simulate. **(G)**. Normalized fraction of calcium ions entering the SA, following a slow calcium entry into the spine head. Here we simulate a single repetition of the injection protocol lasting only 2s showed in Fig. 1H, with *N* =100, 300 and 500 and the pumps removed for *N* =100. Error bars in F & G represent standard errors of the mean (SEM)

To examine the colocalization requirement further, we used our numerical strategy to evaluate how SERCA-ORAI1 distance *d*_SA_ (Fig. 3E), influences the number of ions entering the SA during fast (Fig. 3F) and slow (Fig. 3G) calcium inputs, representing synaptic inputs and SOCE, respectively. Following a fast input with 100 or 300 injected calcium ions we found that the fraction of calcium entering the SA is stable around 15-20% when *d*_SA_ varies from zero to 250nm (Fig. 3F black and red curves). The remaining majority of the ions were found to be extruded by the pumps. To evaluate the influence of the pumps on the calcium uptake, we repeated the same simulations after removing all calcium pumps. In that case, the fraction of SA calcium uptake increases to ≈ 90% (Fig. 3F green curve) as expected, because more ions remain in the cytoplasm.

For the simulation of slow SOCE conditions (Fig. 3G), we placed calcium ions at the top of the spine head with dynamics following the double-exponential fit lasting 2 s with different numbers of ions N. For all the cases (N=100, 300 and 500), the ratio of calcium ions entering SA to the total *N* decreases gradually when *d*_SA_ increases (black, red and blue curves), in contrast to what we observed under the fast calcium input conditions. Additionally, we observed that the distance *d*_SA_ does not influence this ratio of SA calcium uptake when pumps are not present (green). Furthermore, we noted that the fraction of calcium ions entering SA over the total *N* was slightly lower under input conditions when compared to what we saw for the fast inputs. This is likely due to longer RyR activation times during slow calcium inputs (Fig. S7 and S8), resulting in a larger ion loss via diffusion. We also found that the probability of SA calcium depletion via CICR (*P*_2_) is extremely low during a slow calcium input, compared to a fast injection (Fig. S8). In addition, the probability *P*_2_ increased gradually with *d*_SA_, and the time to trigger such events was governed by the presence of calcium pumps in the head.

Therefore, calcium pumps play an unexpected role in preventing and delaying the RyR activation by controlling the arrival of calcium ions at the base of the spine. Taken together, the calcium injection rate, the distance between the SA and plasma membrane, along with the balance of SERCA and calcium pumps shape the SA calcium uptake and the CICR activation probability.

### Calcium dynamics in spines and SA during LTP vs LTD protocols

We next evaluated the consequences of calcium dynamics on the induction of long-term synaptic potentiation and depression (Fig. 4A-C). Although the role of calcium dynamics in LTP and LTD has been known for some time (*8*), the role of SA in these processes remains unclear. It has been previously observed that only the SP+ spines increase their head sizes during LTP (*17*), suggesting that the presence of SA could indeed be a critical factor in the plasticity of the synapse. To examine this possibility further, we used numerical simulations to investigate calcium dynamics during the LTP and LTD induction at a single spine level based on the molecular organization we delineated heretofore.

**Figure 4:**
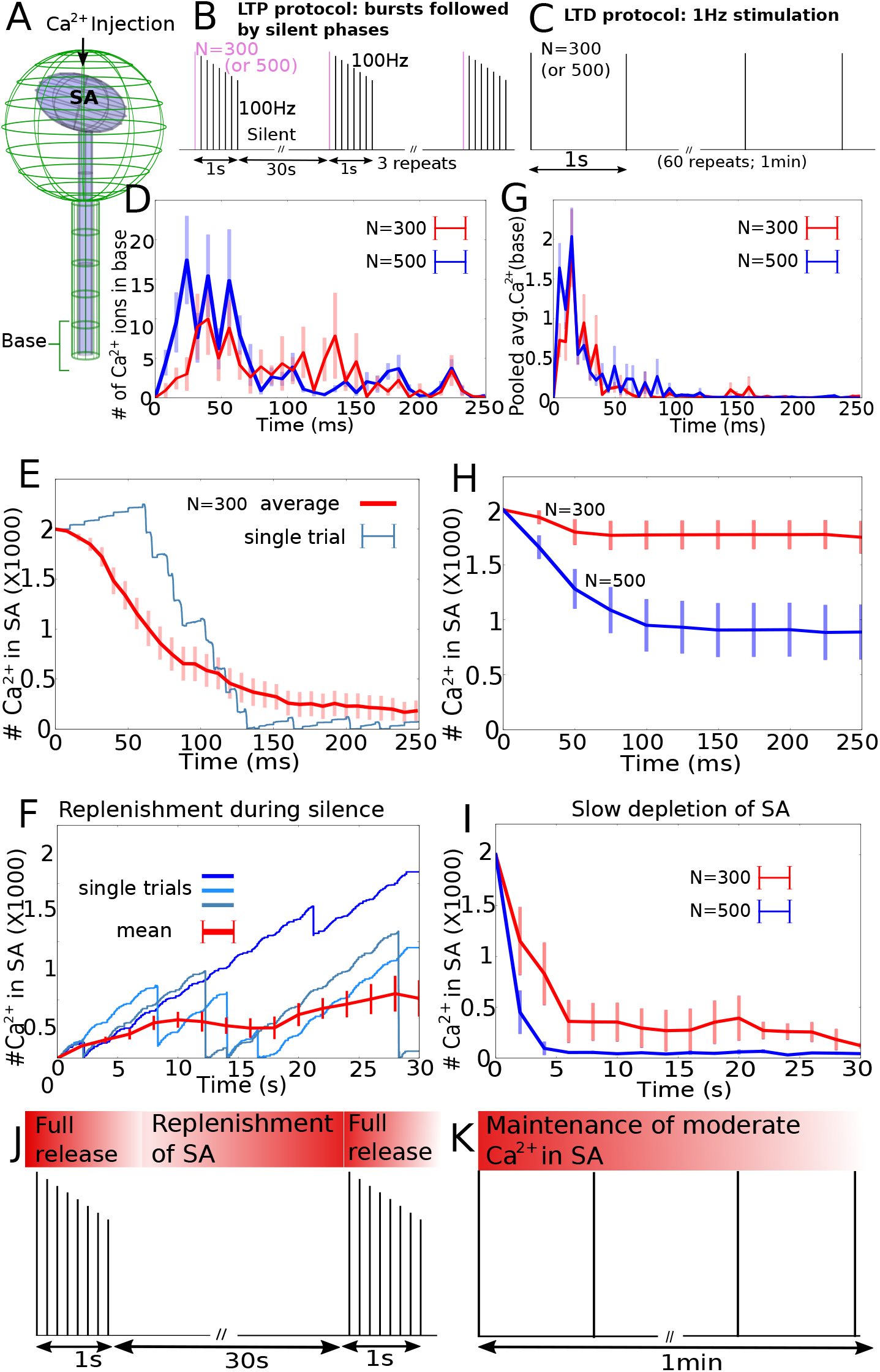
Numerical simulations of calcium transients in a dendritic spine during LTP and LTD induction protocols. **(A)**. Left: scheme of a single spine, showing the SA (light blue) and injected calcium (arrow) in the head. **(B).** Schematic of the postsynaptic LTP protocol with 100Hz stimulations for 1s followed by a 30s silence (SOCE inputs only). The postsynaptic injection rate follows the depression-facilitation model (*26*) (See Methods). **(C).** Postsynaptic LTD protocol is induced by injecting single calcium spikes every 1s and repeated 60 times during 1 minute. We tested the two values *N*=300 & 500 for the number of ions. **(D).** Average number of calcium at the base of the spine during the first 250ms of LTP induction. Error bars show the Standard Error of the Mean (SEM) over 20 realizations. **(E).** Single realization (blue) and average number (red) of calcium ions in the SA within the first 250ms of the LTP protocol (Mean & SEM over 20 trials). **(F).** Simulated SA refilling for the 30s silent phase of LTP. Calcium input into the head is a succession of the slow entry protocol described in Fig. 1H, repeated every 2s. Three typical time courses of the number of calcium ions refilling the SA are shown by the three blue curves, each reaching high, low and medium values at the end of 30s. Mean and SEM values are calculated over 10 trials. **(G).** Averaged calcium response at the base during the first 250ms immediately following each 1Hz spike of the LTD induction protocol. The mean and SEM are calculated over 300 time courses (10 trials and 30s of simulations) for *N*=300 (red) and 500 (blue). **(H).** Number of calcium ions in the SA during the first 250ms of the LTD protocol for *N*=300 (blue) and 500 (black). **(I).** Number of calcium ions in the SA during the LTD protocol shown for 30s for *N*=300 (red) and 500 (blue). **(J).** Interpretation of the LTP protocol as successful CICR at the base followed by SA replenishment. **(K).** Interpretation of the LTD protocol as a maintenance of low calcium in the SA during the 1Hz stimulation.

We simulated spine calcium dynamics during LTP in two phases: (1) stimulation phase and (2) silent phase. The first phase involves a 100Hz calcium spike train, where we injected *N*=300 (or 500) calcium ions. In this range of N, calcium concentration increase falls into the physiological range of ≈0.15*μ*M following a synaptic input (*25*). Afterwards, the injection was allowed to decay slowly with successive events, accounting for synaptic depression. (See Methods for the numerical implementation). Both phases included a slow background input of calcium through the STIM1-ORAI1 pathway (N=300, double exponential timescale lasting 2s, as shown in Fig. 1H). We simulated calcium dynamics for the first 250ms (25 injections) only, and included 30s for the silent phase.

In the stimulation phase, we found that on average, the number of calcium ions in the spine head peaks around 50ms (Fig. S9: blue curve), and then decays with weakened inputs. During this time, less than 10 calcium ions reached the spine base (Fig. 4D); nonetheless they led to several CICR events (green spikes in Fig. S9). Increasing the number of initial calcium ions from 300 (red) to 500 (blue) reduces the onset time of these RyR responses. Due to these depletion events, the stored number of calcium ions in the SA decayed rapidly, leading to a full depletion in about 250ms (Fig. 4E).

During the second phase (that only included slow inputs), we found that on average, a few hundreds of calcium ions replenish the SA through SERCA pumps (Fig. 4F: red curve). In some cases, SA calcium level could go up to more than 1500 ions and sometimes diminish to a very low level due to intermittent CICR events (blue curves in Fig. 4F). We conclude that such variability in SA refilling is compensated by the repetitive procedure of applying high-frequency stimulation followed by silent periods characterizing the LTP protocol. We also repeated the refilling phase of the LTP protocol by adding ectopic vesicular release events that introduce small amplitude calcium spikes (Fig. S10). We found that the presence or absence of ectopic release did not significantly alter SA refilling.

We then investigated calcium dynamics during the LTD protocol (Fig. 4C): calcium ions were injected at a slow rate of 1Hz during one minute. As we confirmed that the transient is much shorter, we decided to simulate calcium dynamics for the first 3 0s. We first injected *N*=300 ions in order to use ion concentrations identical to those employed in the LTP protocol. We then increased the number of ions to *N* =500, and confirmed that results did not change under these conditions. We observed that this input to spine head could trigger CICR events at the base of the spine, that could increase the calcium at very low levels of 10 ions or less when averaged (orange curves in Fig. S11: axis on the right).

In order to compare the LTD responses at the base with the one generated during LTP, we averaged the number of calcium ions during the first 250ms following each 1Hz stimulation pulse (Fig. 4G). The number of calcium ions at the base during LTD simulations is only about one tenth of the LTP simulation response (Fig. 4D). Moreover, the LTP response during the onset of each stimulation lasted longer than the corresponding LTD response. In addition, the calcium depletion in the SA is slower and takes several seconds (Fig. 4H-I), compared to several hundred millisecond timescale observed during the LTP stimulation (Fig. 4D).

We conclude that the SA depletion timescale varies between LTP versus LTD induction protocols, resulting in a strong difference in the calcium levels at the base of the spine (Fig. 4J-K). Overall, we propose that this difference could represent the underlying determinant of the spine’s fate towards either enhancement or depression of synaptic efficacy.

## Discussion

Despite being widely acknowledged as a critical factor of dendritic spine physiology, regulation of the calcium ion dynamics, especially the mechanisms that exert spatio-temporal control over the calcium release and replenishment, remain less clear. We focused on the role of SA in these processes, and discovered that only strong and fast calcium influxes are amplified by the presence of SA in the spine, resulting in CICR at the spine base (Fig. 5A). In contrast, a slow and small amplitude calcium influx through the STIM1-ORAI1 pathway leads to SA replenishment through SERCA pumps located proximal to the plasma membrane (Fig. 5B). From our observations we can conclude that calcium influx timescales, their amplitudes and the spatial distance between ORAI1 and SERCA pumps guarantee that the two pathways (rapid depletion and replenishment) are not triggered at the same time (Fig. 5C).

**Figure 5:**
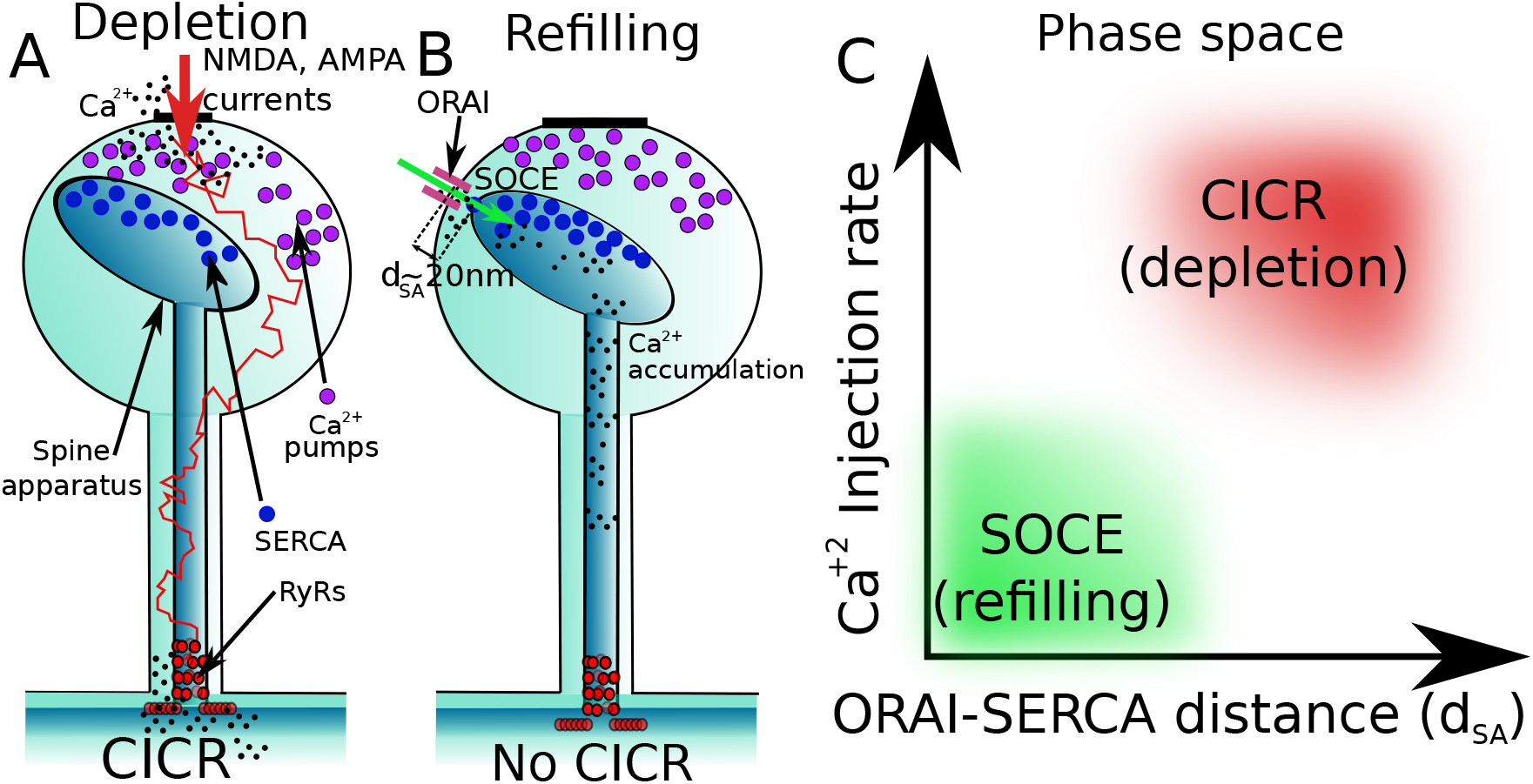
Physiological conditions for SA depletion vs replenishment. **(A)** Synaptic currents entering the dendritic spines through NMDA and AMPA receptors trigger CICR by activating RyRs at the base. **(B)** Small calcium inputs with slow timescales through ORAI1 channels located near the SA membrane are insufficient to trigger a CICR at the base. These ions are either absorbed into calcium pumps located in the spine head, or replenish the SA calcium reservoir via the SOCE through SERCA pumps. **(C)** Phase-space described by the main axes: calcium injection rate and the distance *d*_SA_ between the ORAI1 and SERCA channels. Refilling and depletion conditions of calcium in spines are well separated, so that both do not occur at the same time.

Perhaps the most prominent result of our analysis is the insight that nanoscale molecular architecture plays an essential role in regulating calcium ion release and replenishment. We observed evidence that the distance between the SA and the plasma membrane (*d*_SA_) plays a major role, whereby calcium replenishment stops as *d*_SA_ increases past a certain point. More specifically, we propose that proximity of the two channels ORAI1 and SERCA is required for an optimal SA replenishment, an increase of this distance reduces the chance for the diffusing calcium ions to hit one of the SERCA located in the SA head and thus they could escape to activate RyR located at the base. Interestingly, we did observe that adding more SERCA pumps could partially compensate for a longer distance, suggesting that the effect also depends on SERCA density and distribution. Other key players in these processes are calcium pumps as we show that they restrict small calcium inputs. Indeed, their presence increases the temporal separation of the calcium transient between the head and the base, generating an additional delay in CICR activation.

Although suggested to play in regulating calcium dynamics (*27*), we do not model here calcium buffers because: **(1)** for a fast CICR onset within a 5 ms, an ion bound to a slow buffer would not contribute, thus a simple reduction of the available free ions is equivalent to calcium buffering (*28*). **(2)** In the case of ORAI channel influxes through the plasma membrane, we do not expect buffers to impact SA replenishment due to the short distance *d*_SA_ in the range 20-100nm, which can contain only very few buffers. Nevertheless, similar to surface pumps, buffers could also help separating the depletion from the replenishment regime, as they introduce an asynchrony to the calcium arrival times to the spine base by absorbing calcium ions followed by delayed release. Overall, the present numerical simulations give an upper estimate of the calcium release probability, and therefore, SA replenishment that compensate such depletion events could be possible even with weaker (N<300) SOCE inputs than what we simulated here.

At this stage, we conclude that a few molecular players such as SERCA, RyR and surface pumps seem sufficient to guarantee the interplay between calcium SA refilling and depletion. In the broader context, we propose that these insights could be used to explain the proposed role of the dendritic spines as biochemical computation units (*3*). From this perspective, given that we observe that the SA depletion is achieved only during synaptic inputs and is unlikely during SOCE, the spine can function in an almost deterministic regime.

Main methodological development that enabled our analysis is the construction of the stochastic model that, unlike the models based on average reaction-diffusion equations, which usually ignore molecular details (*25, 29–36*), integrates local binding, and organization of molecules, channels and transporters (*37, 38*). Therefore, the stochastic modeling allows monitoring of diffusion trajectory of each ion separately, thus providing insights at the level of single molecules usually at the expense of high computational cost.

The ability to analyze a system at the single molecule (stochastic) manner is especially important for a system like dendritic spines, given that many of spine processes occur at low copy numbers. For example, AMPARs are of the order of few tens; NMDARs could be less than 10 (*8, 39*). SERCA pumps and RyR are less than hundreds (*40, 41*), and the residual level of calcium in spines also ranges below 100 ions. Here, stochastic modeling allows us to estimate relevant statistics to quantify the fluctuations due to low copy numbers.

Thus in the present modeling and simulations of the spine with a SA, we accounted for the spatial organizations of SERCA, RyR, ORAI1 and SERCA, which allowed us to obtain accurate nanophysiological predictions about the geometrical organization required for replenishment versus release. We were also able to model and examine how calcium ion release and replenishment affects two fundamental processes essential for synaptic plasticity, learning, and memory: LTP and LTD.

Synaptic plasticity is classically thought to result from calcium elevation in spines, activating a variety of molecular pathways. During this process, both spine head volume and the number of AMPARs increase rapidly within a few minutes (*42*). Interestingly, spines with SA predominantly undergo LTP (*15, 17, 43*). Chemical LTP preferentially enlarges and enhances volume of the head and the surface of the post-synaptic density of dendritic spines containing SA (*44*). The need of local calcium increase at the base could lead to protein synthesis, as several machineries such as ribosomes and mitochondria (*45*) are located near the base of the spine neck (*46*). Thus, if local protein synthesis near SA is regulated by calcium ions due to CICR, the products could directly be delivered to the stimulated spines to achieve the postsynaptic changes underlying LTP (*47*) or LTD (*48*).

Our present analysis of calcium dynamics during the LTP-LTD protocols revealed the store depletion level, the total duration, the overall strength and the initial height of the calcium response at the spine base. At this stage, it remains unclear whether there is a molecular mechanism that compares calcium transients between spine head and base during synaptic plasticity. However, we showed the number of calcium ions at the base during LTD is only about 10% of the calcium concentration during LTP. This increased calcium concentration at the base during potentiation has been previously shown to regulate protocols is in agreement with the observations that trafficking of such as AMPA and or NMDA receptors (*8, 49–51*). It would therefore be interesting to investigate how these receptors could be selected by local calcium elevation. Additionally, this elevation at the base could also restrict receptor diffusion and interactions, result in potential well nanodomain trapping (*52, 53*) or phase separation (*54*), and create an asymmetric receptor influx in spines. Finally, calcium elevation at the base of spines could trigger ER-mitochondria calcium communication (*45, 55*) to produce ATP required for the spine homeostasis and spine shape and volume remodeling (*56*).

We also identified a key difference in SA depletion timescales between LTP and LTD, suggesting that this is also a key determinant of whether a given dendritic spine enhances or depresses synaptic signal. Moreover, in our simulations of the LTD protocol, failure to induce CICR prevents SA depletion and leads to a larger calcium accumulation. Thereby subsequent inputs could trigger a large calcium increase at the base of the spine. This scenario could explain the instability of LTD induction that could accidentally result in LTP, especially in the presence of a secondary calcium source such as voltage-gated channels.

Although these additional aspects of calcium dynamics in dendritic spines remain to be examined in future studies, our current work discovered the critical role that molecular architecture plays in regulating calcium ion release and replenishment in dendritic spines. The architecture imposes constraints on the two opposing processes, thus ensuring fidelity and spatio-temporal control which ensures that calcium stores within SA are replenished without triggering calcium release.

## Materials and Methods

### Ethics statement

Animal handling was done in accordance with the guidelines of the Institutional Animal Care and Use Committee (IACUC) of the Weizmann Institute of Science (Approval number: 00650120-3 from 20/1/2020 for 3 years), College de France and the appropriate Israeli and French laws and national guidelines. Experiments were carried out according to the guidelines of the European Community Council Directives of January 1, 2013 (2010/63/EU) and of the local animal welfare committee. All efforts were made to minimize the number of used animals and their suffering.

### Calcium imaging experiments

#### Culture preparation

Cultures were prepared as detailed elsewhere (*57*). Briefly, rat pups were decapitated on the day of birth (P_0_), and their brains were removed and placed in a chilled oxygenated Leibovitz L15 medium (Gibco) enriched with 0.6% glucose and gentamicin (Sigma; 20 *μ*g/ml). Hippocampal tissue was dissociated after incubation with trypsin and DNAase, and passed to the plating medium consisting of 5% horse serum and 5% fetal calf serum prepared in minimum essential medium (MEM, Gibco), enriched with 0.6% glucose, gentamicin and 2 mM GlutaMax (Gibco). Approximately 10 000 cells in 1 ml of medium have been plated in each well of a 24-well plate, onto a glial layer which has been grown for a week before the plating of the neurons. Cells were left to grow in the incubator at 37°C, 5% CO2 for 3 d, following which the medium was switched to 10% horse serum in enriched MEM, and in addition of 5′fluoro-2-deoxyuridine + uridine (FUDR) (Sigma; 20 *μ*g and 50 *μ*g/ml, respectively), to block glial proliferation. The medium was replaced 4 d later by 10% horse serum in MEM. The same medium was used after the transfection and no further changes were made until cultures were used for experimentation.

#### Transfection

Transfection was conducted at 7-8 day in vitro (DIV). A Lipofectamine 2000 (Invitrogen) mix was prepared at 1 *μ*l/well with 50 *μ*l/well OptiMEM (Invitrogen), and incubated for 5 min at room temperature in the hood. Separately, a mix of 2 *μ*g/well total DNA in 50 *μ*l/well OptiMEM was prepared and also incubated for 5 min. Then, two preparations were co-incubated for 15 min at room temperature in the hood. This mix was then added to the each culture well at the amount of 100 *μ*l/well, and allowed to induce the transfection during 3 h before a final change of medium. In most cases, at least 20 neurons/well were transfected. In these experiments SP-short subcloned into pEGFP-C1 (BD Biosciences, Clontech) (*58*) or into mCherry were used. For morphological analysis, a blue fluorescent protein (BFP) plasmid was cotransfected with the SP construct. Co-transfected cells displayed no apparent differences in spontaneous calcium activity, morphology, spine density and survival compared with BFP-only transfected cells or non-transfected cells. The distribution and pattern of the expression of the SP plasmid were similar to those of the endogenous SP (*12*). Co-transfection efficiency for the plasmids using this method is nearly 90%. Experiments were conducted routinely at 7-10 days after transfection. Cultures were used at the same age for comparisons.

#### Imaging and drug application

Cultures were incubated for 1 h in Fluo-2 (high affinity) AM (2 *μ*m; Invitrogen, Carlsbad, CA, USA) containing recording medium containing (in mM) NaCl 129, KCl 4, MgCl_2_ 1, CaCl_2_ 2, glucose 4.2, and HEPES 10; pH was adjusted to 7.4 with NaOH, and osmolality to 320 mOsm with sucrose. Alternatively, K+ Fluo-4 salt solution was injected into transfected neurons with sharp micropipettes and allowed to diffuse for 0.5 hours before imaging. In the latter case, no BFP transfection was required and cell morphology could be detected based on Fluo-4 basal fluorescence. After loading of calcium sensor cells were imaged using LSM 880 Zeiss (Germany) upright confocal microscope equipped with 40x 1 NA water-immersion objective. Spontaneous calcium transients were detected in both SP and BFP co-transfected and non-transfected neurons using fast scan mode (10-20 Hz / frame). Bath application of the following blockers: tetrodotoxin (TTX, 1 *μ*M), APV (50 *μ*M) and DNQX (20 *μ*M) (all from Sigma) has been used to eliminate action potentials, PSPs, neurotransmitter release and activity-induced calcium transients. Caffeine (5 mM) was added using quick bath perfusion and then washed out rapidly, while fast calcium transients from dendritic segments with SP(+) and SP(-) spines of co-transfected cells were recorded. In some cases, a caffeine-containing patch pipette (diameter 1-2 *μm*, 15 mM) was positioned close to individually identified dendritic segments of a SP-transfected neuron and responses to local pressure application of caffeine in neighboring SP+ and SP-spines of the same dendritic segment were imaged. Images were obtained at high speed for detecting rapid changes in [Ca2+]i (10-20 Hz, restricted, horizontally oriented scan region). In latter case, distance between the caffeine containing pipette and individual dendritic spines was carefully similar for all cases.

#### Data analysis

Fluorescence intensity was calculated using ZEN (Zeiss, Germany), ImageJ (NIH, Bethesda, MD, USA) and Matlab software (MathWorks Inc., Natick, MA, USA). Dendritic protrusions were categorized into spine types based on their morphological measurements. Two independent observers conducted some of the analysis. SP+ and SP- dendritic spines that were used for calcium imaging were identified in BFP-transfected of Fluo-4 microinjected neurons, and analysed independently. Statistical comparisons were made with t-tests or ANOVA, as appropriate, using Matlab and KaleidaGraph (Synergy Software, Reading, PA, USA).

### Experiments using STED

#### AAV production and injection

This was performed as previously described in (*16*). Briefly, a GFP cassette was placed under the control of a hSynapsin promoter in a serotype 9 AAV virus. Two-month old C57Bl6 mice were anesthetized under ketamine/xylazine in 0.9% NaCl. AAVs were diluted in PBS to 1.02×1013 vg/mL and 1 *μ*L of virus was injected into the hippocampal region. After two weeks, the mice were scraficied and the brains extracted after 2% PFA/PBS intracardiac perfusion.

#### Immunohistochemistry and STED microscopy

This also performed as previously described in (*16*). Briefly, 40-*μ*m thick brain slices were permeabilized and blocked for 2 h in 0.25% Triton/0.2% Gelatine in PBS at room temperature. Primary and secondary antibodies were diluted in the same solution, incubated 2 h at room temperature followed by overnight at 4°C. In addition to the primary and secondary antibodies used in (*16*), anti-ORAI1 [Mouse, 1:100, Abcam ab244352], anti-Synaptopodin (SP) [Guinea Pig, 1:100, Synaptic System 163004], anti-chicken Star Green [Goat, 1:200, Abberior STGREEN-1005] and anti-guinea pig Alexa 405 [Goat, 1:500, Abcam, ab175678] were used in this study. For the distance measurements between SERCA3 and ORAI1, Z-stacks of 100 nm steps images were taken using a 3-color super-resolution 3D-STED microscope (Abberior Instruments GmbH, previously described in (*16*)). Note that the distances that are measured below 100nm are located in the same Z plane, as the Z step of our stack acquisition is 100nm. Thus, the errors in the distance measurements are minimized compared to 2D microscopy systems.

Super resolution was used for SERCA3 and ORAI1 channels. All channels were then deconvolved using Huygens software and analyzed using an in-house developed plugin on ImageJ to measure the distance between two maximas in 3D (maximum distance of 400 nm as a cutoff threshold). Between 83 and 120 interactions (distance from ORAI1 to SERCA3 less than 400 nm) per slice were analyzed in the spine head, between 27 and 52 in the neck and between 45 and 88 at the base. Four slices were analyzed. The violin plots represent all the dots analysed (378 in the head, 159 in the neck and 290 at the base).

For the presence of SP, images were taken with another set of immunostained slices with an additional 405 nm laser for the excitation of the SP-alexa405, using a Zeiss Axioobserver Z1 with a CSUW11 Spinning-Disk scan head (Yokogawa 63x/1.4NA objective lens). Z-stacks of 150 nm steps were taken and analyzed with ImageJ software. Colocalization frequencies were calculated over the total number of spine head analysed. 19 field of views from 4 animals were analysed. 60 to 80 spine heads were analysed per field of view. Each interaction type was assigned to one of the five categories of colocalization type.

#### Statistical analysis

All data are expressed as mean ± SEM. Statistical significance for within-group comparisons was determined by one-way ANOVAs (followed by Tukey’s post-test) on GraphPad Prism.

### Stochastic model of calcium dynamics and numerical simulations

#### Modeling and simulation of calcium diffusion

##### Stochastic simulations of calcium ions in a dendritic spine

To simulate calcium transients in a spine, we use the following model: the dendritic spine geometry is made of a large spherical head connecting the dendrite by a cylindrical neck (*21, 59*). We added a spine apparatus as a “spine inside a spine” with a similar geometry (Fig. 2D). The parameter for radii of the spine head, spine neck, SA head and SA neck are summarize in Table S2.

The motion of calcium ions are modeled with Brownian diffusion described by the stochastic equation 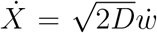, where *w* represents Wiener white noise, delta-correlated in both space and time: for distinct time and space coordinates *X, X′* and *t, t′* therefore, 〈*w*(*X, t*)*w*(*X′, t′*)〉 = *δ*(*X − X′*)*δ*(*t − t′*), where *δ*(.) is Dirac’s delta function. We simulate a discretized form of this motion using the Euler’s scheme: 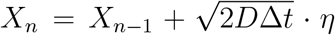. Here *X_t_* = {*X, Y, Z*} is the position of a particle at time *t* and η is a normal random variable with three independent components generated by the NumPy library of Python. The diffusion coefficient of calcium in the medium is *D* while Δ*t* is the width of a single time step (values in Table S2). We chose the largest Δ*t* such that reducing it further neither alters the calcium fluxes through SERCA pumps nor the RyR activation times.

We consider the small baseline concentration of free initial calcium in the medium to be zero, thus we introduce calcium in two ways: either we position instantaneously a total of *N* calcium ions at single point at the top of the spine head (fast synaptic inputs) or we introduce calcium ions one after the other according to a distribution which follows a difference of two exponentials (STIM-ORAI pathway: described below).

After entering inside the spine, ions can diffuse within the spine head until it reaches the bottom of the neck. Spine base is modeled as an absorbing boundary, thus ions arriving at the base of the spine do not appear again in the simulation. In our model, we neglected any electrostatic interactions between the ions and the membranes of the spine or the SA, from which ions are reflected with the classical Snell-Descartes law.

Moreover, calcium ions have two valence charges which can create an electric interaction with the surface charge density located on the dendrite membrane. However, in the presence of a small calcium influx, the possible interaction of the ion with the rest of the medium is described by the Debye length in an order of a few nanometers (*60, 61*). Therefore, as there is a minimum distance of 20nm between the two membranes in our model, we neglected the electro-diffusion of calcium ions.

##### Calcium extrusion pumps

Calcium extrusion pumps are located on the inner surface of the spine head, and are modeled as absorbing circular disks having a catchment radius of 10nm as previously calibrated in (*59*). To match a decay time scale of 6ms recorded for calcium fluorescents in the spine head, we calibrated the number of pumps to be 50 (*16*). Indeed, if we increase this catchment area, we would need to reduce the number of pumps to keep this calcium decay time fixed.

##### Ryanodine receptors

We model RyRs as circular disks located on the surface of SA with a catchment radius of 10nm where calcium ions are bound. There are *n_R_*=36 RyRs located at the base (Fig. 2D). Out of these, 12 are located on the segment of the SA parallel to the dendrite. The remaining 24 form four rings (six receptors in each ring) in the SA neck. Opening of a RyR is triggered by the arrival of two calcium ions into the receptor site. When a first calcium ion arrives at a receptor, it stays bound for 10ms, then unbinds to diffuse to a distance of one Brownian step. The RyR is opened only if the second one arrives within this 10ms window. We confirmed (not shown here) that results in RyR opening times and probabilities are largely independent of this window size, when varied from 10ms to. After the arrival of a second ion to the RyR calcium ions are released with a delay of 0.25 ms. The number of calcium ions released per RyR starts from 8 and decays to 6 and 7, followed by another cycle of 8-7-6 as reported in (*16*). After each release, RyRs are inactivated for 3 ms, during which they do not bind to calcium ions.

#### SERCA Pumps

Classical models of SERCA pumps are based on a four-state Markov chain model, where most of the parameters are unknown (p.43, after equation 2.47 in (*36*)). We model here SERCA pumps using the same formulation as we previously implemented in (*16*) with a stochastic model of four states: 0 ions bound, 1 ion bound, 2 ions bound & refractory state. A pump is opened by the arrival of two successive ions within a 10ms window. If a second ion does not arrive within this time, the first ion is released at a distance given by one Brownian step. In case of an opening event, the two ions gets translocated into the SA and the pump remains inactive for its refractory period. Hence our model uses only two parameters: the first ion’s waiting time of 10ms and the refractory time of 100ms. The catchment radius of SERCA pumps R=10nm is justified by the atomic-level description given in (*62*). The positioning of SERCA pumps are on the top hemisphere of the SA head, according to a uniform random distribution (Fig. 2D).

#### Influx through the STIM1-ORAI1 pathway

We do not model here explicitly the transfer of ions from the extracellular to the intracellular medium upon the activation of ORAI1 channels that form complexes with STIM molecules. Instead, to simulate the SOCE inputs through ORAI channels, we use the time course extracted from the calcium fluorescence signal of the spine head (Fig. 1C and Fig. S 3). In addition, we vary the distance *d*_SA_ between the plasma membrane and the SA membrane that governs the proximity between ORAI1 channels and SERCA pumps.

### Simulation of LTP and LTD protocols for calcium injection

1. The 1s high frequency stimulation during the LTP protocol is simulated here as a series of calcium spikes into the top of the spine head. The amplitude of this spike is a decreasing number of ions proportional to the fraction *E* of synaptic resources in the effective state we compute using the facilitation-depression model (*26*) described by:

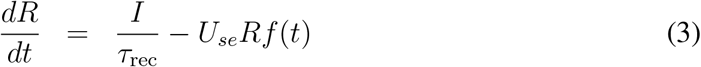

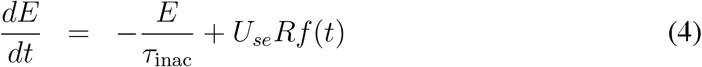

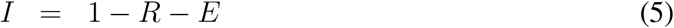

Here *I* and and *R* are the inactive and recovered fractions of synaptic resources that add up to the normalised amount of total resources with value 1. The two time constants *τ*_rec_=0.3s and *τ*_inac_=0.2s govern the recovery and the inactivation of the resources, respectively (*63*). The stimulation protocol is m d by the function *f* (*t*) made of 100Hz train of *δ* Dirac impulses during 1s: 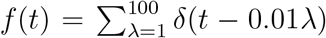. Finally, *U_se_* is the fraction of synaptic resources in the recovered state getting activated by each instantaneous input. During the first 250ms we simulated, the value of *E* decayed from 1 to 0.47, which depends weakly on *U_se_*. This decay indeed simulates the gradual reduction of the synaptic input amplitude into spines. In addition to these injections of fast calcium spikes, we also maintain the slow input of calcium (through ORAI1 channels as in Fig. 1F with *N*=300 ions and the long 2s duration). During the simulation of the refilling phase of the LTP protocol (Fig. 1F), the high-frequency stimulation is absent, hence, only this slow calcium influx was available.
2. For each stimulation pulse of LTD, we injected calcium ions instantaneously every 1s over a duration of one minute (we only simulate the first 30s of the protocol to investigate the SA calcium dynamics). We pre-determined the number of ions contained in injected pulse to be *N*=300 or 500. In addition to these fast injections, throughout all LTP simulations we also inject a repetition of the slow calcium influx as before.

## Author contributions

DH, EK, NR and KB designed the study. KB and DH conducted the mathematical analysis. KB implemented the computational simulations. EK, DM, LK performed the experiments and analysed the data. AB provided the virus construction. NR, DM and EK edited the manuscript.

